# AdaGenes: A streaming processor for high-throughput variant annotation and filtering

**DOI:** 10.1101/2025.06.17.659929

**Authors:** Nadine S. Kurz, Klara Drofenik, Kevin Kornrumpf, Kirsten Reuter-Jessen, Jürgen Dönitz

## Abstract

**Motivation:** The amount of sequencing data resulting from whole exome and whole genome sequencing (WES / WGS) presents challenges for annotation, filtering, and analysis. These challenges are exacerbated by the need for efficient and scalable tools that can handle the vast amounts of data generated by modern sequencing technologies.

**Results:** We introduce the Adaptive Genes processor (AdaGenes), a sequence variant streaming processor designed to efficiently annotate, filter, LiftOver and transform large-scale VCF files. AdaGenes provides a unified solution for researchers to streamline VCF processing workflows and address common challenges in genomic data processing, e.g. to filter out non-relevant variants to focus on further processing of the relevant positions. AdaGenes integrates genomic, transcript and protein data annotations, while maintaining scalability and performance for high-throughput workflows. Leveraging a streaming architecture, AdaGenes processes variant data incrementally, enabling high-performance on large files due to low memory consumption. The interactive front end provides the user with the ability to dynamically filter variants based on user-defined criteria. It allows researchers and clinicians to efficiently analyze large genomic datasets, facilitating variant interpretation in diverse genomics applications, such as population studies, clinical diagnostics, and precision medicine. AdaGenes is able to parse and convert multiple file formats while preserving metadata, and provides a report of the changes made to the variant file.

**Availability and implementation:** A public instance of AdaGenes is available at https://mtb.bioinf.med.uni-goettingen.de/adagenes. The source code is available at https://gitlab.gwdg.de/MedBioinf/mtb/adagenes.

## 1 Introduction

The exponential growth of next-generation sequencing (NGS) data presents new opportunities for identifying disease-causing genetic variants. However, the sheer scale and complexity of these datasets increasingly challenge variant interpretation workflows. A critical bottleneck lies in efficiently reducing millions of raw variants to a focused set of pathogenic or oncogenic variants to investigate the origins of genome-related diseases. While variant annotation and filtering tools transform variant data into actionable insights, current solutions remain fragmented: Command-line and web-based annotation pipelines [1, 2, 3, 4, 5], offer robust variant processing, but demand computational expertise for deployment and customization. Second, there are VCF (Variant Call Format) filtering tools [6, 7, 8, 9, 10, 11, 12, 13, 14, 15, 16, 17] that enable criterion-based prioritization, such as population allele frequency or ClinVar reports, yet operate independently from annotation steps. In addition, command-line based tools require experience in setting up and defining filtering rules manually on the command line. Complementary visualization tools [18, 19, 20] provide variant exploration and analysis of VCF files, but often lack integrated processing capabilities. Furthermore, with HGVS nomenclature gaining prominence, normalization and validation tools [21, 22] address syntax challenges, but remain siloed from variant interpretation pipelines. This fragmentation urges researchers to manually chain multiple tools - each with distinct installation, runtime, and syntax requirements - creating workflow inefficiencies and accessibility barriers.

Here we present a fast and powerful tool, the Adaptive Genes processor (AdaGenes), for variant annotation as well as filtering genetic variant data enhanced by a versatile web interface. AdaGenes is capable of processing whole exome sequencing (WES) variant files and provides a solution for LiftOver, variant annotation, sorting, filtering, data transformations, HGVS-compliant normalization and interactive variant data exploration. By unifying these capabilities into a single platform, Adagenes accelerates WES analysis and the discovery of pathogenic variants in biological and clinical research.

## 2 Materials and methods

### 2.1 The AdaGenes server

The AdaGenes server was implemented using Python (v3.11) and Flask (v2.2.5). To reduce the memory capacity required to process WES and WGS files, variants are read in batches of 5000 variants each, which are processed in parallel. Further, the batches are split into chunks of 100 variants, which are sent to the annotation modules and merged after all annotation threads are completed. Given a variant set *𝒱*, the batches *ℬ*, the chunk size *𝒞*, the thread per batch process *T*, and the set of annotation processes *𝒜*, we can define the variants as a set of batches, where the per-batch parallel chunk processing is formalized as the parallel processing of the annotations, which are then assembled again.

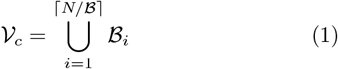

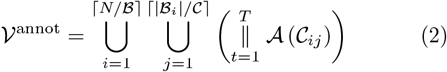

### 2.2 The AdaGenes database

The database storing the uploaded variant data is a MongoDB database, which enables the storage of dynamic, previously unknown annotation features due to its NoSQL approach [24]. For each uploaded file, AdaGenes creates a new collection identified with a unique ID for the uploaded variants, as well as a second collection for storing metadata, including the genome version, added annotations, and VCF column definitions. Each variant is assigned an HGVS identifier based on the standardized nomenclature for genetic variants of the Human Genome Variety Society (HGVS) [29], the order of the variants is ensured by saving the line number within the original file. We can define the variant database *𝒟* as a set of collections *C*, including a set of variants *𝒱c*, where each variant entry *v* is a set of key value pairs of the available features *K* and possible values *U*

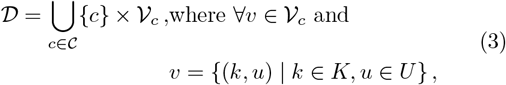

In case of a LiftOver, AdaGenes adds the positions of other reference genomes as annotations. After each annotation, an index is created for each new annotation features, enabling fast variant filtering based on the annotation features.

### 2.3 The AdaGenes web front end

The AdaGenes web front end was built using Vue.js (v3). Layout formatting was implemented using the Vuetify (v3) library. The interactive table for displaying sequence variants and filtering options was implemented using AG Grid. In order to make the front end usable for large variant files, status queries were implemented in addition to the main data query, which regularly check the current status of data processing. To display large variant files, including whole exome and genome sequencing, we only load a small number of variants from the backend at a time. Interactive visualizations were implemented using Plotly [25]. All modules were containerized using Docker [23].

### 2.4 Data preprocessing

To enable the processing of different input data formats, we implemented multiple custom data parsers for AdaGenes, including variant data parsers for VCF, MAF, BED, CSV and XLSX files. To process these different file formats, AdaGenes automatically identifies the file type and employs the associated reader..gz-compressed files are extracted using the gzip package. Each file reader parses the data into an internal, JSON-based data structure, which allows further processing independent of the data structure of the original file. Similarly, we implemented various writers being able to export variants in different file formats, including VCF, MAF, CSV, and JSON. Liftover between genome assemblies was implemented using chain files of the UCSC FTP server [26] and the Python LiftOver package.

### 2.5 Variant annotation

To annotate variant data, Onkopus [27] was integrated for annotating variants. Genomic locations were mapped to transcript and protein level using SeqCAT [28]. To annotate variants with pathogenicity scores, the Onkopus module of the dbNSFP database [30] was queried. In the case of multiple values, the highest pathogenicity score was selected. In addition, the dbNSFP rank scores were added to the annotation, scaling the predictions of each method to the 0 to 1 scale to allow comparability of the methods. As some annotation modules require pre-annotations of other modules, including the gene association of protein-coding variants, the annotation process has been divided in two stages. Within these stages, requests to the Onkopus modules were parallelized to increase the annotation performance.

### 2.6 Transcript and protein to genome conversion

To convert variant data at transcript or protein level to genome level, we query the SeqCAT Application Programming Interface (API) based on the hgvs package [31], and extract the chromosome, position, as well as reference and alternate alleles. Given that the transcript information is not contained at the protein level, protein identifiers are mapped to the MANE Select transcript [32] of the respective gene.

### 2.7 Benchmarking datasets

Multiple publicly available variant files have been used for testing AdaGenes and generating the visualizations, including datasets on single nucleotide variants, insertions and deletions, and structural variants [33, 34, 35, 36, 37, 38] (Table S1, available as supplementary data).

## 3 Results

### 3.1 AdaGenes provides comprehensive variant annotation

We present AdaGenes, a high-throughput streaming processor for sequence and structural variant annotation and filtering. Its versatile web interface provides a straight-forward application that allows the annotation, filtering and data transformations of genetic variant data into various formats. AdaGenes employs a three-tier architecture: The server, the database, and the web front end (Fig. 1). The input file may be a variant file in VCF or MAF format, or a custom table-formatted format (e.g. CSV). When a file is uploaded, AdaGenes converts the variant data into an internal, JSON-based representation and stores the variant data in a NoSQL database. The AdaGenes server can perform 4 different actions: It can 1) convert table-formatted data from custom file formats to VCF, 2) perform LiftOver between different reference genomes, 3) annotate variants, and 4) sort and filter variants (Fig. 1). The server is able to perform different annotation and filtering steps, including the conversion of variant data at transcript and protein to genomic level, the LiftOver between genome assemblies (GRCh37, GRCh38, T2T-CHM13), annotation of variants, as well as sorting and filtering of variants.

**Figure 1.**
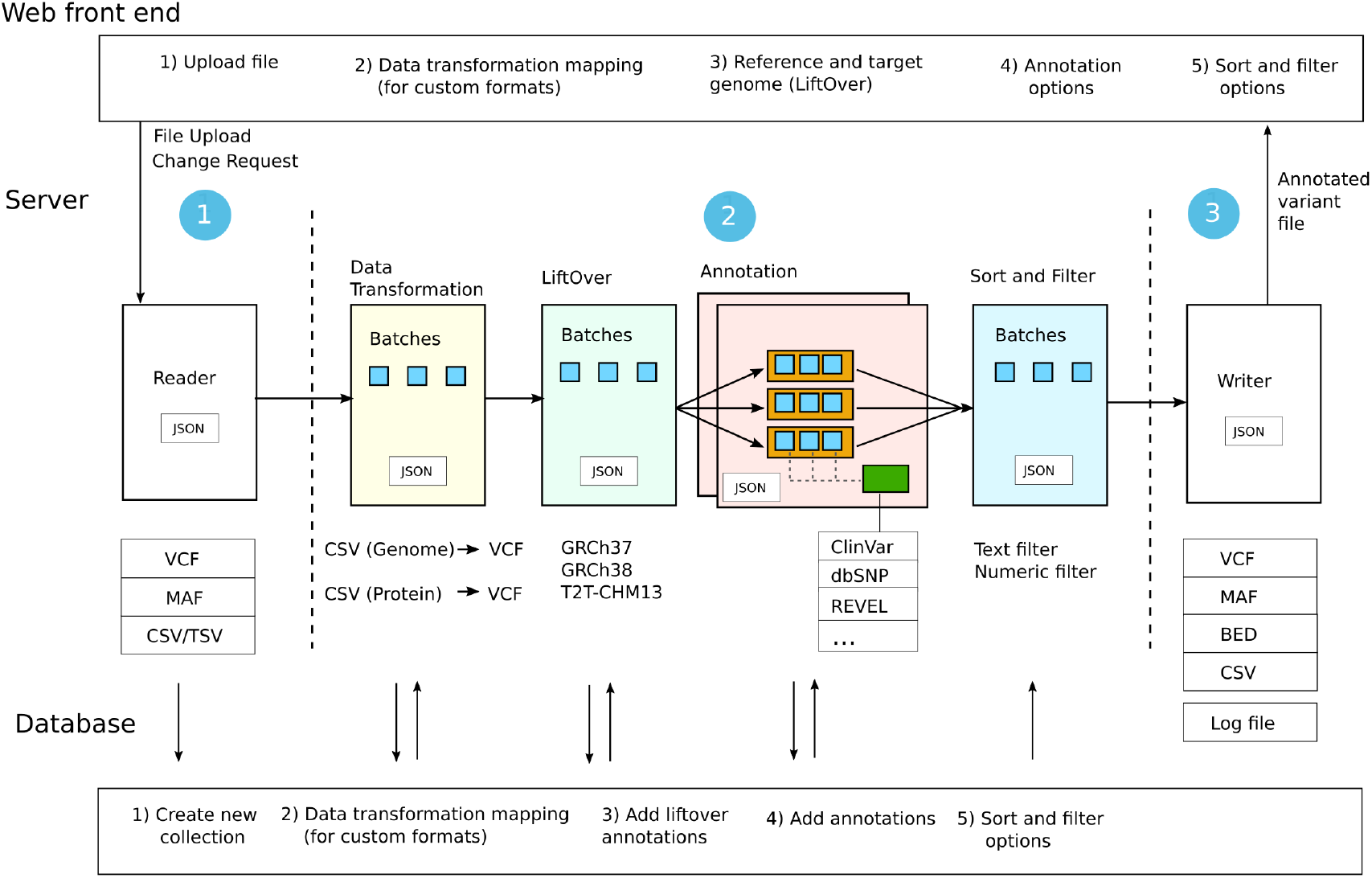
The AdaGenes server workflow. The AdaGenes server reads a variant file and converts it into an internal JSON-based representation (1). The variants are stored in a NoSQL database, creating a new collection for an uploaded variant file. AdaGenes supports 4 kinds of variant processing steps: The conversion of table-formatted data (e.g. CSV) at transcript or protein level to VCF, LiftOver between reference genomes, the annotation of variants, and the sorting and filtering of variants (2). In all modules, large amounts of data are efficiently processed in small data blocks (batches, blue boxes). To accelerate annotation, the batches are divided into smaller parts (chunks, red boxes), which are sent to the annotation modules (green box). The processed variants can then be written out again in one of the supported file formats, as well as a log file of all made modifications (3).

We provide a comprehensive annotation of variants for AdaGenes, including genomic, transcript and protein data, population allele frequency, pathogenicity prediction scores, functional regions, ClinVar reports, and gene expression. In addition, AdaGenes provides multiple novel annotation features related to the molecular and structure-specific characteristics of variants, protein sequences, and drug-gene interactions (Table S2, available as supplementary data). We compute multiple molecular characteristics of the reference and alternate amino acid, including the change in molecular weight, charge, hydropathy, flexibility, solubility, or ionization and phosphorylation potential. Structure-specific variant features are derived based on AlphaFold-predicted protein structures of the MANE Select transcripts, and include surface accessibility, secondary protein structure, Phi and Psi angles, as well as hydrogen bonds at the mutated site. For annotation, the data is divided into smaller batches and annotated in parallel across multiple CPU cores. In order to annotate variants in the web front end, the user uploads a variant file in a standard format (VCF, MAF, CSV) and selects the corresponding reference genome (Fig. 2a (1)). By selecting a different target genome, AdaGenes automatically performs a LiftOver to the selected target genome, while preserving the former genomic positions as annotations. In order to present variant annotations in a readily legible format, we generate an interactive table that comprises the predefined columns of the VCF format - CHROM, POS, ID, REF, ALT, QUAL and FILTER - as well as a separate column for each annotation feature (Fig. 2a (2)). The user can now sort and filter the variant data, or annotate it with additional features. The annotation is possible by selecting one, multiple, or all modules for consultation from a list of available annotation modules (Fig. 2a (3)). The annotation time per module depends on the database size and the complexity of the queries. The estimated expected processing speed can be viewed by clicking on the tool tips that are displayed next to each annotation module. Once the annotation process is complete, the variant table is automatically updated and the newly annotated properties are displayed as additional columns. After each processing step, such as LiftOver, annotation, sorting, and filtering, the results can be downloaded in various file formats, including VCF, MAF and CSV (Fig. 2b). We generate a unique ID for each uploaded document, providing persistent links to view the analysis results again later.

**Figure 2.**
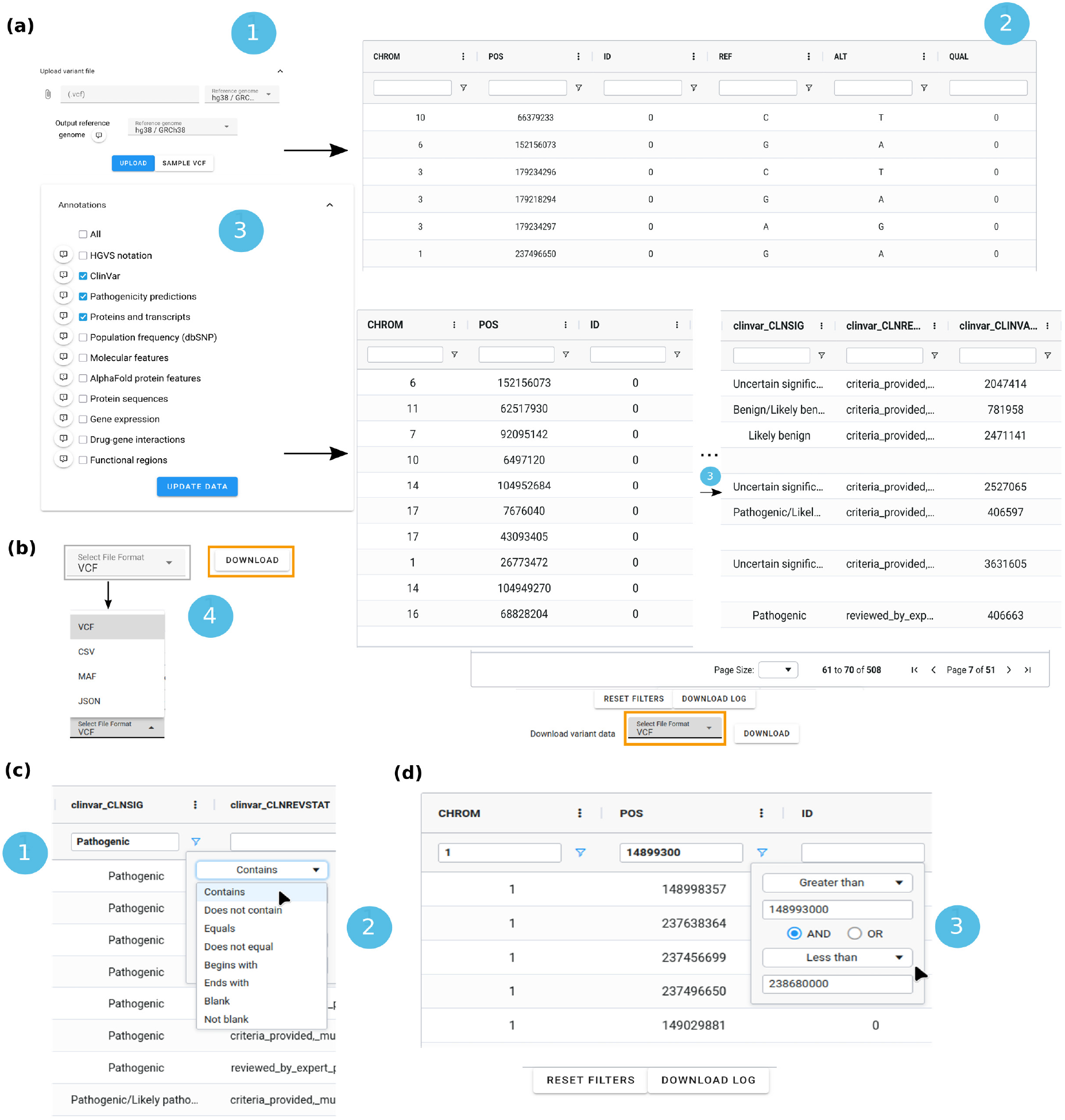
Interactive annotation using AdaGenes. (a) In addition to the file to be uploaded, the reference genome of the variant data and the target reference genome can be specified (1). AdaGenes then generates an interactive table in the selected reference genome, where the VCF columns as well as each INFO annotation feature is a separate column. After uploading the file, one or multiple annotation modules can be selected (3). After the annotation process, new features are added as columns that can be sorted and filtered (3). (b) The annotated variants can be downloaded multiple formats, including VCF, MAF, CSV and JSON (4). (c) Feature-specific variant filtering options in AdaGenes. Variants can be filtered (1) with a free text search, (2) refined by defining a query operator, including text-specific filtering options, as well as (d) with the generation of complex numeric queries (3) using AND/OR operators.

### 3.2 AdaGenes enables dynamic interactive variant filtering and sorting

Following the annotation of variants, they can be sorted and filtered directly according to the newly added features. After a file has been uploaded, AdaGenes identifies the data type of each column based on the VCF metadata or the column data entries and automatically generates corresponding filtering options. Filters can be created using a text input form (Fig. 2c(1)) and refined by selecting a query operator (Fig. 2c(2)). For numeric features, complex queries can be created by concatenating filters using AND/OR operators (Fig. 2d(3)). Table 1 presents a comprehensive overview of the main available functions of AdaGenes, alongside a comparison with other variant annotation and filtering tools. All filter and sorting options can also be reset directly via a button in order to carry out a new filtering process. To ensure reproducibility, AdaGenes also provides a downloadable log file, which records all modifications made to the original file, including annotations, LiftOver and applied filters.

**Table 1.**
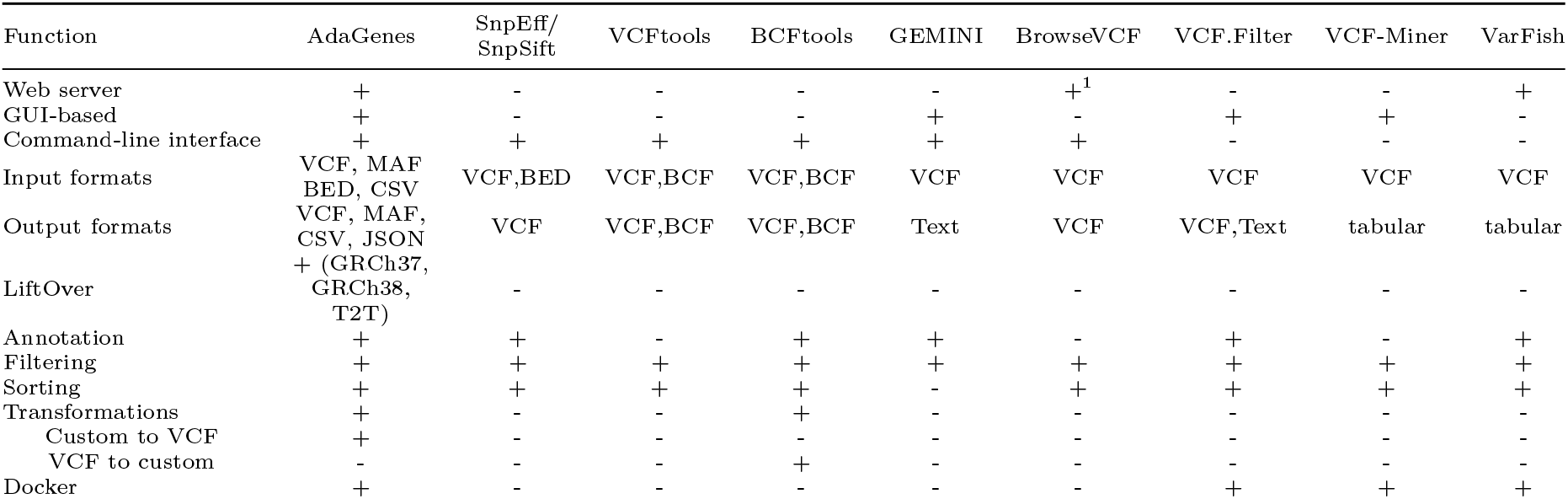
Comparison of AdaGenes to other variant annotation and filtering tools.

### 3.3 Conversion from transcript and protein level to genomic data

In addition to data from sequencing pipelines, biomedical research frequently includes datasets on genetic variation data available in proprietary data formats. For example, the CIViC database [39] is a resource for the expert-curated data on the interpretation of cancer variants. Single nucleotide variants that have been identified as biomarkers are referred to at protein level (e.g. ‘KRAS G12C’), making it difficult to annotate or analyze the variants with tools that require standardized data formats at the genomic level. For this purpose, AdaGenes provides the functionality to convert custom table-formatted data files into VCF format. In case an uploaded file does not correspond to the known genomic variant data formats, the data is displayed in its original form, and the transformations options are enabled (Fig. 3(2)). To convert the data from transcript and protein to genome level, users can select the respective columns in their data that correspond to the gene names and amino acid exchange (Fig. 3(3)).

**Figure 3.**
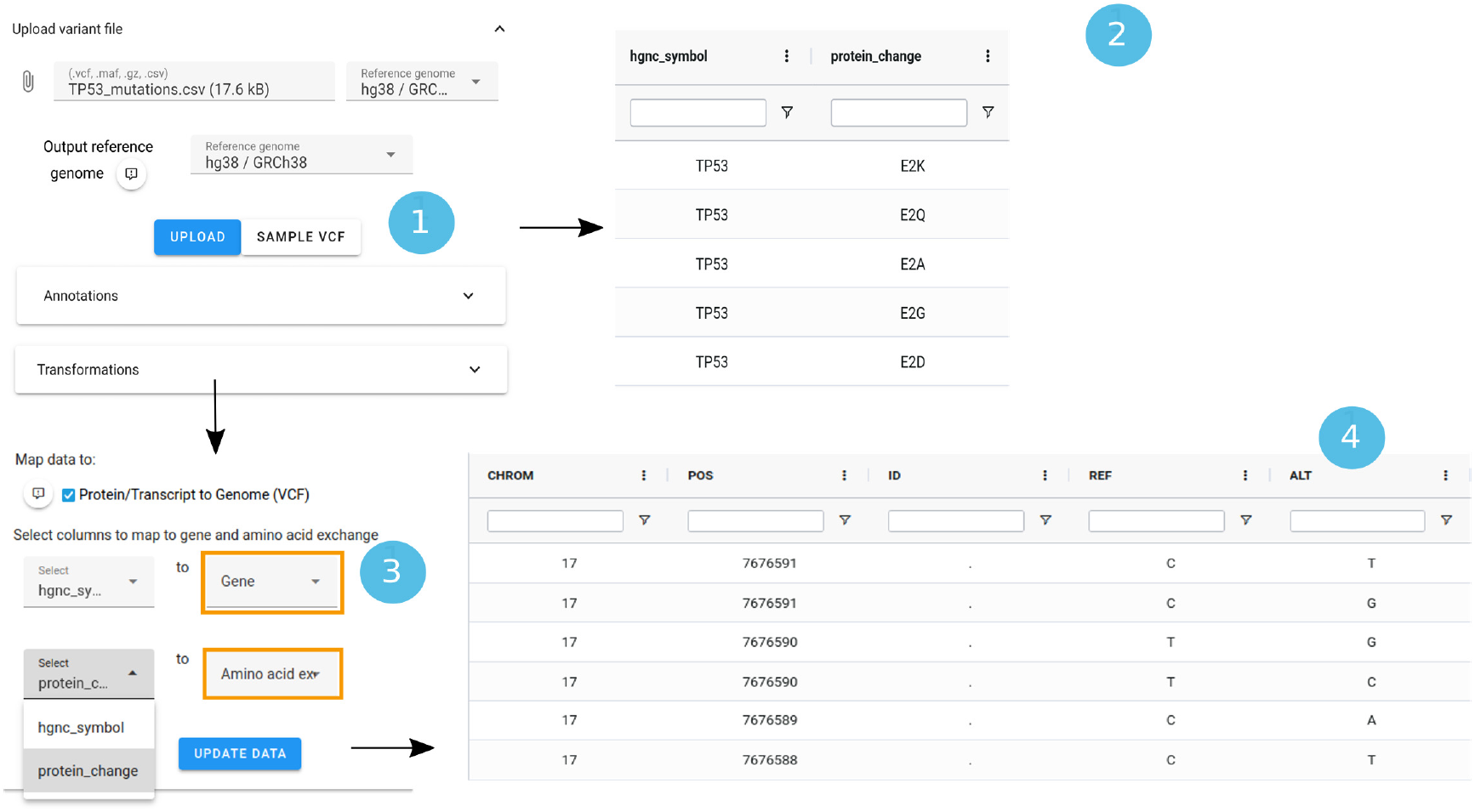
Protein to genome conversion. Users can map custom data formats at transcript or protein level to VCF format by uploading a file in a custom data format (1). Recognizing a file format different from commonly known formats, AdaGenes displays the custom data (2) and enables the data transformations form (3). Users can the select the columns in their data representing the gene names and amino acid exchanges. AdaGenes then converts the variants from protein to genome level and generates a VCF representation (4).

### 3.4 AdaGenes provides dynamic statistics and analysis of molecular profiles

The AdaGenes server computes statistics and interactive visualizations of the processed variant data, which are generated dynamically in real-time and adapted to the selected filters. including the frequency of the different mutation types (Fig. 4a), and the 20 most pathogenic variants (Fig. 4b). By applying filters, the variants can be analyzed dynamically in more detail. For example, selecting a specific gene would allow to examine the pathogenicity of the variants within that gene (Fig. 4c). The analysis of the number of variants per gene can be refined by adding a ClinVar filter, to view only the pathogenic variants within the affected genes of a molecular profile (Fig. 4d). In addition, a molecular feature filter could be added to filter the variants within mutated protein domains (Fig. 4e).

**Figure 4.**
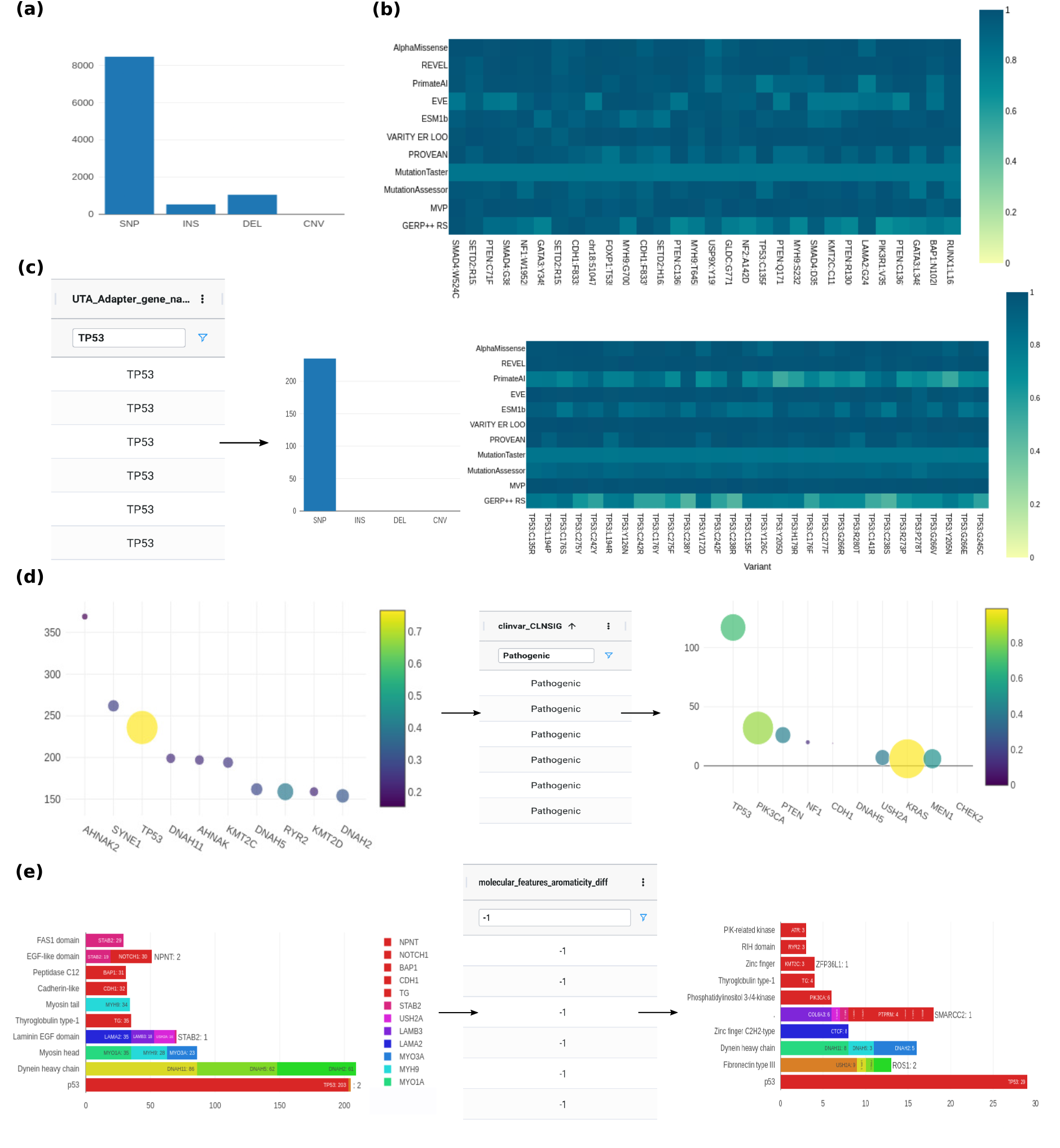
Interactive molecular profile analysis. (a) By interactively filtering variants in the web front end, variants can be dynamically analyzed according to different criteria, including the number of variant types in the uploaded variant files, and (b) the most pathogenic variants according to several pathogenicity prediction scores (0=benign, 1=pathogenic). To refine the analysis of the pathogenic variants of the dataset, filtering options can be applied, e.g. the selection of a specific gene (c), to further narrow down the variant selection. (d) Representation of the number and pathogenicity of variants in different genes. Here, all variants within a gene are counted, where the circle size represents the average AlphaMissense-predicted pathogenicity of all variants within each gene. By selecting one or multiple filtering criteria, e.g. a ClinVar entry, the variants in genes can be analyzed according to specific characteristics, like the investigation of pathogenic variants only. (e) Domain-specific hotspot mutation analysis, featuring the number of variants found in protein domains in different genes. By adding a molecular filter, like the selection of variants with a loss of aromaticity, users can analyze the biochemical processes of genetic variants at the molecular level.

### 3.5 AdaGenes provides a dual-use concept comprising a web application and a command-line interface

Featuring a command line interface (CLI) and a web front end, AdaGenes offers two modes for annotating variants: For the command-line based usage, input files are split into several files, which are annotated in parallel, and merged again after the annotation has been finished. In addition to the web application, which focuses on user-friendliness and interactivity, the command line tool enables the professional generation of annotation pipelines.

In addition, it is possible to install AdaGenes locally and use it in environments with restricted Internet access, e.g. clinical infrastructures. To perform the annotation completely locally, e.g. for sensitive patient data, Onkopus and SeqCAT can also be installed locally.

## 4 Discussion

We presented AdaGenes, a modular streaming processor for annotating and filtering genetic variant data. Compared to existing tools, AdaGenes combines all steps necessary to preprocess and filter molecular profiles, including LiftOver, annotation, filtering, and variant analysis in one tool. AdaGenes’ two-tier parallelization strategy for variant annotation capture both memory efficiency and computational parallelism. The modular architecture enables the seamless expansion or update of annotation modules without the need to stop the database and the server.

In comparison to other annotation pipelines, AdaGenes provides a broader range of annotations and uniquely offers the ability to annotate molecular and protein structure specific features of protein-coding variants. AdaGenes’ structural protein feature predictions (e.g., hydrogen bonding changes, surface accessibility) enable novel insights into variant pathogenicity mechanisms—particularly valuable for missense variants in cancer or rare diseases. The capacity to import a variety of data formats, including VCF, MAF and BED, and to export to other data formats, offers a broad spectrum of potential applications, ranging from clinical to biomedical research. The ability to generate filter options based on previously unknown annotation features makes the tool more dynamic and versatile than previous solutions with fixed filter options. The option of defining filters in the graphical user interface provides a more intuitive filtering of variants than the manual definition of filters via command-line-based tools. In addition, AdaGenes is a versatile tool for visual exploration and analysis of variant data. An innovative feature is the computation of dynamic analytics. where visualizations and statistics adapt in real-time to user-defined filters, facilitating rapid hypothesis testing and data refinement - a feature absent in preliminary tools. In this way, AdaGenes enables the analysis of the biochemical characteristics of molecular profiles, helping in exploring the molecular processes behind variant pathogenicity. The recording of all modifications and the option to export them as a log file ensures the reproducibility of all results. The web server is free to use, requires no login or registration and does not transfer data to third party partners. The AdaGenes server and web front end as well as the annotation modules, including Onkopus and SeqCAT, are locally installable and can thus be utilized in clinical environments to process sensitive patient data, e.g. in molecular tumor boards. AdaGenes is thus ideal for use in precision medicine to annotate and pre-filter WES and WGS files. The remaining subset of pathogenic variants can be exported and subsequently loaded into a tool for detailed variant analysis and targeted therapy selection, e.g. Onkopus. We are thus confident that AdaGenes provides a platform that can significantly enhance the accessibility of genomic analysis, empowering more researchers and clinicians to interpret variants accurately and drive innovative biological insights.

## 5 Acknowledgments

We thank Tim Tucholski and the molecular tumor board of the University Medical Center Göttingen for their support and valuable feedback. We also thank the International Max Planck Research School for Genome Science (IMPRS-GS) for their support. We acknowledge support by the Open Access Publication Funds of the Göttingen University.

## 6 Author contributions

JD and NSK conceptualized the architecture and method. The software was implemented by NSK (Database, server, web front end) and KD (Annotation parallelization). KK optimized the annotation process. KRJ contributed test cases and evaluated the methodology. NSK wrote the manuscript. All authors proofread and approved the manuscript.

## 7 Supplementary data

Supplementary data are available at Bioinformatics online.

